# Automated Analysis of Medial Gastrocnemius Muscle-Tendon Junction Displacements During Isolated Contractions and Walking Using Deep Neural Networks

**DOI:** 10.1101/2020.09.29.317529

**Authors:** Rebecca L. Krupenevich, Callum J. Funk, Jason R. Franz

## Abstract

Direct measurement of muscle-tendon junction (MTJ) position is important for understanding dynamic tendon behavior and muscle-tendon interaction in healthy and pathological populations. Traditionally, obtaining MTJ position during functional activities is accomplished by manually tracking the position of the MTJ in cine B-mode ultrasound images – a laborious and time-consuming process. Recent advances in deep learning have facilitated the availability of user-friendly open-source software packages for automated tracking. However, these software packages were originally intended for animal pose estimation and have not been widely tested on ultrasound images. Therefore, the purpose of this paper was to evaluate the efficacy of deep neural networks to accurately track medial gastrocnemius MTJ positions in cine B-mode ultrasound images across tasks spanning controlled loading during isolated contractions to physiological loading during treadmill walking. Cine B-mode ultrasound images of the medial gastrocnemius MTJ were collected from 15 subjects (6M/9F, 23 yr, 71.9 kg, 1.8 m) during treadmill walking at 1.25 m/s and during maximal voluntary isometric plantarflexor contractions (MVICs). Five deep neural networks were trained using 480 labeled images collected during walking, and were then used to predict MTJ position in images from novel subjects 1) during walking (novel-subject), and 2) during MVICs (novel-condition). We found an average mean absolute error of 1.26±1.30 mm and 2.61±3.31 mm in the novel-subject and novel-condition evaluations, respectively. We believe this approach to MTJ position tracking is an accessible and time-saving solution, with broad applications for many fields, such as rehabilitation or clinical diagnostics.

## Introduction

Ultrasound imaging is an increasingly popular tool for quantifying *in vivo* muscle-tendon dynamics in humans, with fidelity to do so during functional activities such as walking and running (1, 2). Tracking distinct anatomical landmarks in recorded ultrasound images - namely, the muscle-tendon junction (MTJ) - allows researchers the ability to differentiate muscle from tendon behavior with applications in clinical gait analysis (3), rehabilitation from injury (4, 5), and efficacy of exercise interventions (6). Traditionally, analyzing cine B-mode ultrasound images requires the researcher to manually label landmarks of interest within each frame in a video sequence. This process is time consuming and labor intensive (e.g., a single 2 s video recorded at 70 frames/s would require labeling 140 frames) and is thereby not practical for larger datasets across multiple cohorts with many experimental manipulations or conditions. In recent years, several semi- or fully-automated tracking tools have been developed for the purpose of tracking and quantifying muscle fascicle length and orientation (7–9), but these approaches are generally ineffective at tracking MTJ displacement. Thus, there is a need for alternative solutions to accurately and efficiently track the MTJ during functional activities.

Deep learning approaches to image processing have been successful in medical image analysis (10) and may be a promising technique for automated MTJ tracking. Recent advancements in deep neural networks (DNNs), such as the use of convolutional layers, have improved performance across a wide range of feature detection tasks (11, 12). DNN-based architectures consist of multiple hidden layers (i.e. deep layers) that ‘learn’ common features of the training images, such as shading or shapes, and then use that learned knowledge to recognize those features in future images(13). The addition of deep convolutional and deconvolutional layers allows DNNs to detect learned features anywhere in the image, which makes this approach particularly well-suited for analysis of medical images (e.g., ultrasound). Leitner et al. (2020) recently demonstrated that deep learning with a convolutional neural network can successfully track the MTJ during isolated maximal contractions and passive rotation (14). However, to our knowledge, these methods have not been evaluated on ultrasound images recorded during more functional activities with physiological loading such as walking. Further, it is unknown the extent to which a DNN that was trained on images from one movement type can be generalized to tracking a novel movement.

Widespread adoption of deep-learning methods for image processing in the biomechanics community has been limited by the availability of accessible solutions for researchers without extensive experience in deep learning. User-friendly open-source software packages, such as DeepLabCut (15, 16), have removed this barrier to deep learning methods but their efficacy has not yet been demonstrated for tracking MTJ displacements in cine B-mode ultrasound images. Therefore, the purpose of this paper was to evaluate the efficacy of deep neural networks constructed with DeepLabCut (15, 16) at accurately tracking medial gastrocnemius MTJ positions in cine B-mode ultrasound images across tasks spanning controlled loading during isolated contractions to physiological loading during treadmill walking. We aim to benchmark the performance of these analytical techniques against conventional manual tracking approaches. Trained networks presented in this paper will be made publicly available with the peer-reviewed manuscript.

## Methods

### Subjects

15 subjects (6M/9F, 23 yr, 71.9 kg, 1.8 m) participated in this study. Prior to participation, subjects were screened and excluded if they reported injury or fracture to the lower-extremity within the previous six months, neurological disorders affecting the lower-extremity, or currently taking medications that cause dizziness. All subjects provided written informed consent according to the University of North Carolina Biomedical Sciences Institutional Review Board.

### Dataset

Subjects first walked for six minutes at 1.25 m/s on an instrumented treadmill to pre-condition their triceps surae and allow their movement patterns to stabilize prior to data collection (17). Subjects then walked for two minutes at 1.25 m/s (Bertec, Columbus, Ohio) while a 10 MHz, 60 mm linear array ultrasound transducer (LV7.5/60/128Z-2, Telemed Echo Blaster 128, Lithuania) operating at 38-68 frames/s recorded cine B-mode images of the gastrocnemius MTJ. Three subjects were collected at 38 frames/s due to an unanticipated change to the settings file, the remaining 12 subjects were collected with frame rates between 60-76 frames/s – the later range of frame rates reflects differences in window sizes (e.g 80% vs 100% window) between subjects. One 10-s video was recorded during the last 15 seconds of the two-minute walking trial. Two consecutive full strides of walking were identified and isolated using GRF data to determine the timing of gait events. On a separate day, subjects performed two maximal voluntary isometric plantarflexor contractions (MVIC) while seated in a dynamometer (Biodex Medical Systems, Shirley, NY) while we recorded cine B-mode images of the gastrocnemius MTJ. For each subject, we selected one video and trimmed the video so that it begins with the subject in a relaxed state and ends when the subject reaches peak torque (corresponding to greatest MTJ displacement). For all subjects, the same investigator manually labeled the gastrocnemius MTJ in each frame of the isolated video with respect to the distal image boundary, the results of which we interpreted as our ground truth measurement. The position of the MTJ was identified as the most distal insertion of the muscle into the free tendon (Fig. 1A). The full subject set of walking videos was then randomly split into five different combinations of training set (n=12) and test set (n=3), such that each subject was represented in the test set in one of the five combinations. To account for between-subject differences in stance time and framerates, 80 frames were randomly chosen from each subject (1200 frames total) for use in training and evaluation of the networks. The MVIC videos were set aside for use in a later testing stage (novel condition testing), as described below, and were not used to train the networks.

**Figure 1.**
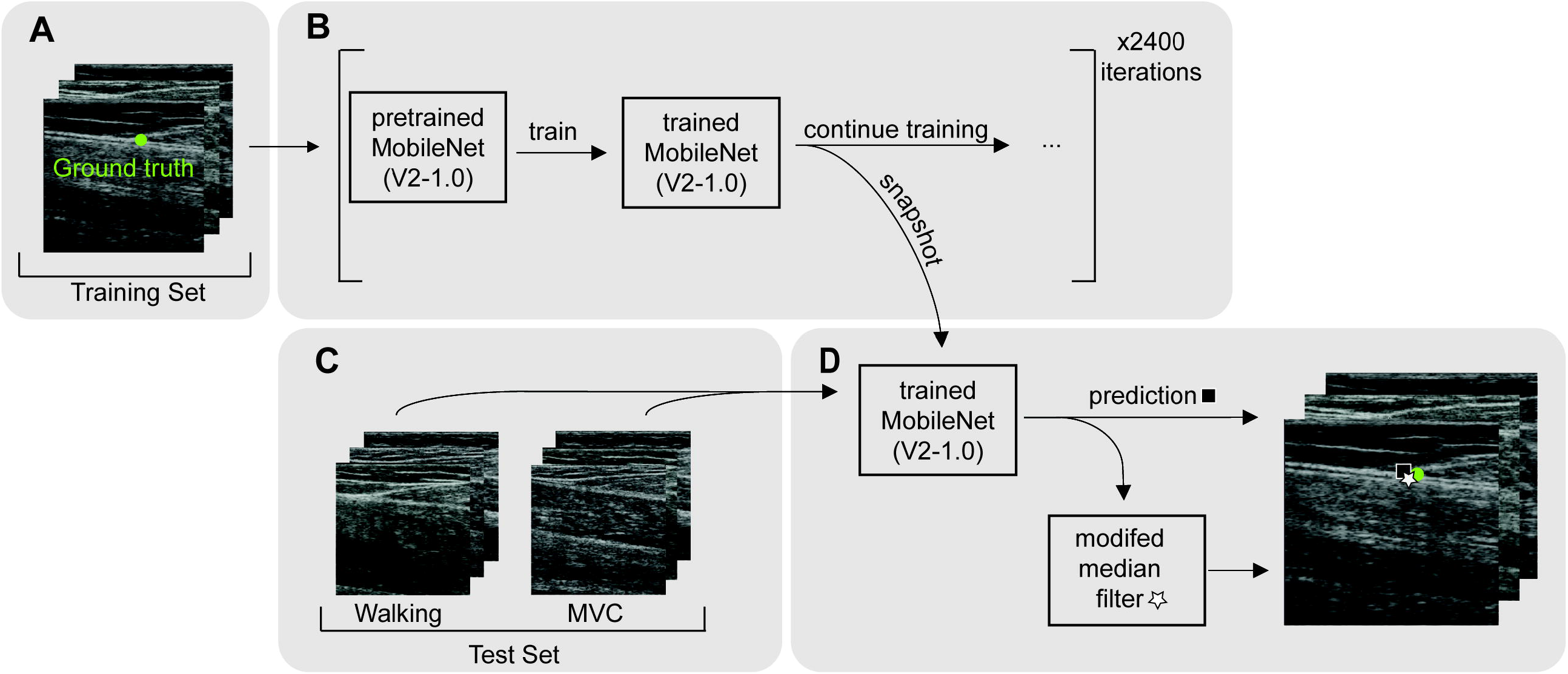
A) MTJ position was manually labeled (ground truth) for each frame of B-mode ultrasound video collected during walking. Five training sets were created, consisting of 80 frames from 12 subjects. B) Networks were based on MobileNetV2-1.0, pretrained on ImageNet with default parameters, and then trained on our training sets for 24000 iterations. During training, the networks were saved intermittently (i.e. a snapshot was taken) to allow for evaluating the effect of training time on performance. C) Each training set was associated with a test set consisting of three subjects that were not included in the respective test set (i.e., novel subjects). D) The trained networks were used to predict the location of the MTJ in each frame of the videos from the novel subjects in the test set. A modified median filter was applied to predictions with low confidence scores (<0.98) to reduce noise from outlier data points.

### Network

We used DeepLabCut (Version 2.1.6.4) (15) to construct and train our networks. Networks were based on MobileNetV2-1.0, pretrained on ImageNet (18) with default parameters, which were then trained on our training sets for 18,000 iterations with a batch size of 20. We systematically varied the size of the training set and trained 45 distinct networks (five each of 80, 60, 40, 20, 10, 5, 2, and 1 frame(s) per subject included in the training sets). During training, the networks were saved intermittently to allow for evaluating the effect of training time on performance.

### Evaluation

After training, each network was used to predict the location of the MTJ in each frame of the videos from each of the three subjects in the network’s respective test set. Each prediction is accompanied by a confidence score (0-1) which represents the probability that the MTJ is visible in the frame (15). To reduce noise from outlier data points, we applied a modified median filter to predictions with low confidence scores (<0.98). The default window size was seven frames, such that a low confidence prediction at frame *i* is replaced with the median of the set of predictions from frames *i* – 3 to *i* + 3. This window size was reduced at the beginning and end of videos to maintain symmetry around the target frame. For example, at frame *i* = 2 and *i* = 3, the window size is reduced to three (frames 1-3) and five (frames 1-5) frames respectively. We report both unfiltered and filtered data in the results.

#### Novel Subject Evaluation

Each network was used to track MTJ location in each frame of the novel subjects (i.e. completely independent from the training set) from its respective test set, as detailed above (Fig 1B). From these frames, 80 were randomly chosen from each subject for evaluation.

#### Novel Condition Evaluation

To further characterize generalizability, the networks were used to track cine B-mode images of the gastrocnemius MTJ collected during maximal voluntary isometric contractions (MVIC). One subject was excluded from this analysis due to data loss (n=14). Each network was evaluated with 40 randomly chosen frames from each of the novel subjects from its respective test set. Novel condition images were not used in any of the training sets.

Network performance was quantified using root mean square error (RMSE) and mean absolute error (MAE). RMSE was calculated for the Euclidean distance between the predicted and ground truth MTJ positions. MAE was calculated as the Euclidean distance between the predicted and ground truth MTJ positions. RMSE and MAE were calculated for each individual subject. We also report overall RMSE, which represents the RMSE of all subjects, and average MAE, calculated as the group average MAE. Finally, we report the percentage of valid or invalid predictions. A valid prediction was defined as being within a 5 mm radius of the ground truth (14) (Fig. 2).

**Figure 2.**
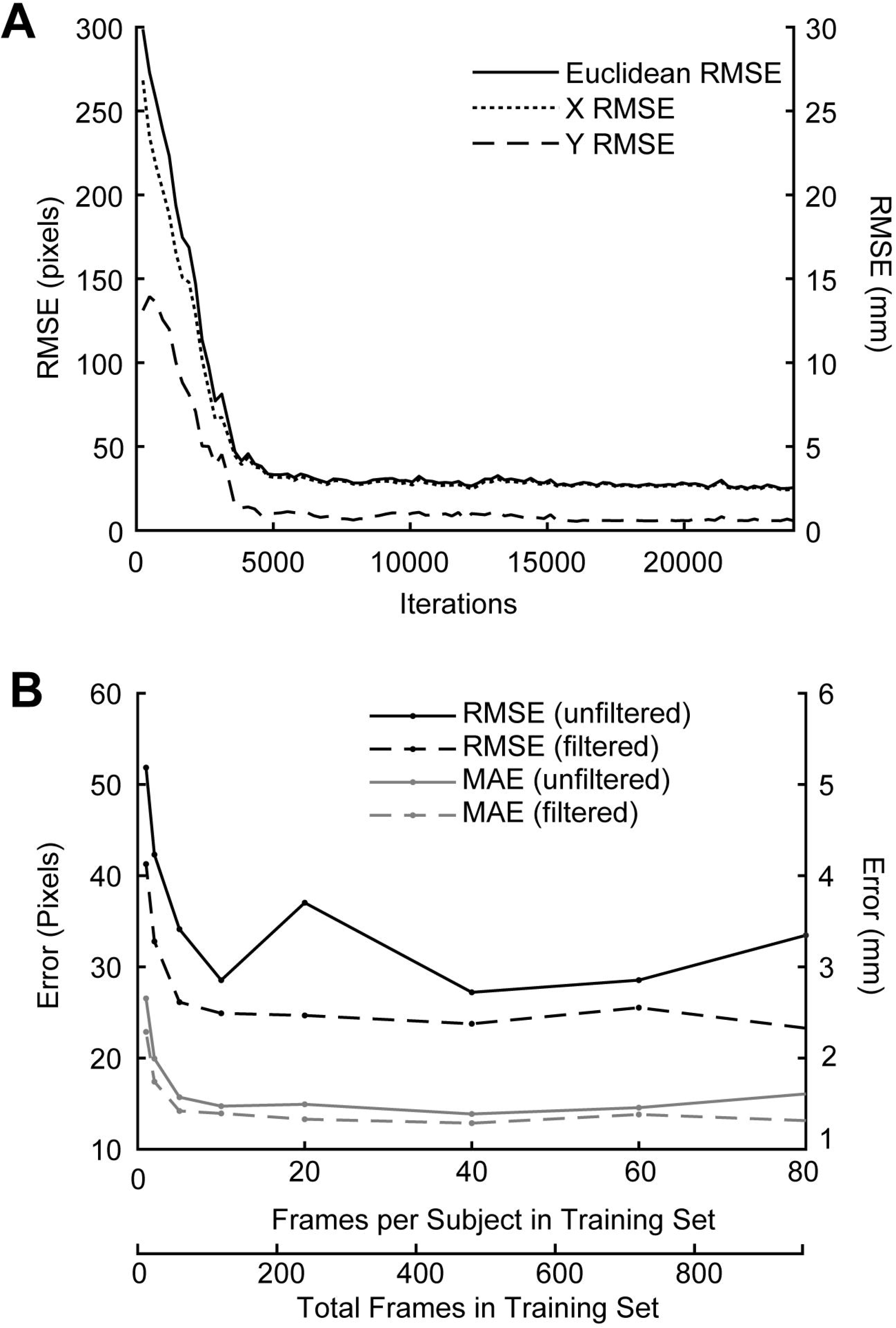
A) Root mean square error (RMSE) between ground truth and predicted MTJ position was evaluated every 5,000 iterations during training. RMSE showed a steep initial drop and subsequent plateau around 4,000 iterations, and was minimized at 18,000 iterations. B) RMSE and MAE decreased with an increasing number of frames, but was relatively stable from 500 to 1000 frames. 480 labeled frames (40 frames per subject) were sufficient to achieve an overall RMSE of less than 3 mm and an overall MAE less than 2 mm. RMSE is reported in pixels (left axis) and mm (right axis).

### Speed Benchmarking

We evaluated GPU inference speed on Nvidia Tesla V100-SXM2 16 GB and Nvidia GeForce GTX1080 with a batch size of 64. CPU inference speed was evaluated on a 2.5 GHz Intel Xeon (E5-2680 v3) processor with batch sizes of 64 and 8. Average inference speed (frames/s) is reported for the novel-subject evaluation.

## Results

### Training parameters

After a steep initial drop at ~3,000 training iterations, RMSE fell below 5 mm and remained stable as the number of iterations increased. RMSE was minimized at ~18,000 training iterations. (Fig. 2A). For training set size (i.e. total number of frames in training set), we found that RMSE and MAE decreased with a greater number of frames and were both relatively stable from 500 to 1000 frames. A training set size of 480 total frames (40 frames per subject) was sufficient to achieve an overall RMSE of less than 3 mm and an overall MAE less than 2 mm (Fig. 2B).

### Novel-subject evaluation: medial gastrocnemius MTJ displacements during walking

Overall unfiltered RMSE in the novel-subject evaluation was 2.72 mm, with an average unfiltered MAE of 1.26±1.30 mm (Table 1). The modified median filter reduced overall RMSE by 0.35 mm (2.37 mm) and MAE by 0.09 mm (1.17±1.23 mm). 94 and 95% of unfiltered and filtered predicted novel-subject MTJ positions were classified as valid (i.e., ≤ 5 mm of ground truth) (Table 2). MAE for three subjects in the novel-subject evaluation was greater than two standard deviations from the mean (Fig. 3A). Removing these subjects resulted in a smaller overall RMSE (Unfiltered: 1.36 mm; Filtered: 1.11 mm) and a smaller average MAE (Unfiltered: 0.70±0.49 mm; Filtered:0.65±0.35 mm).

**Figure 3.**
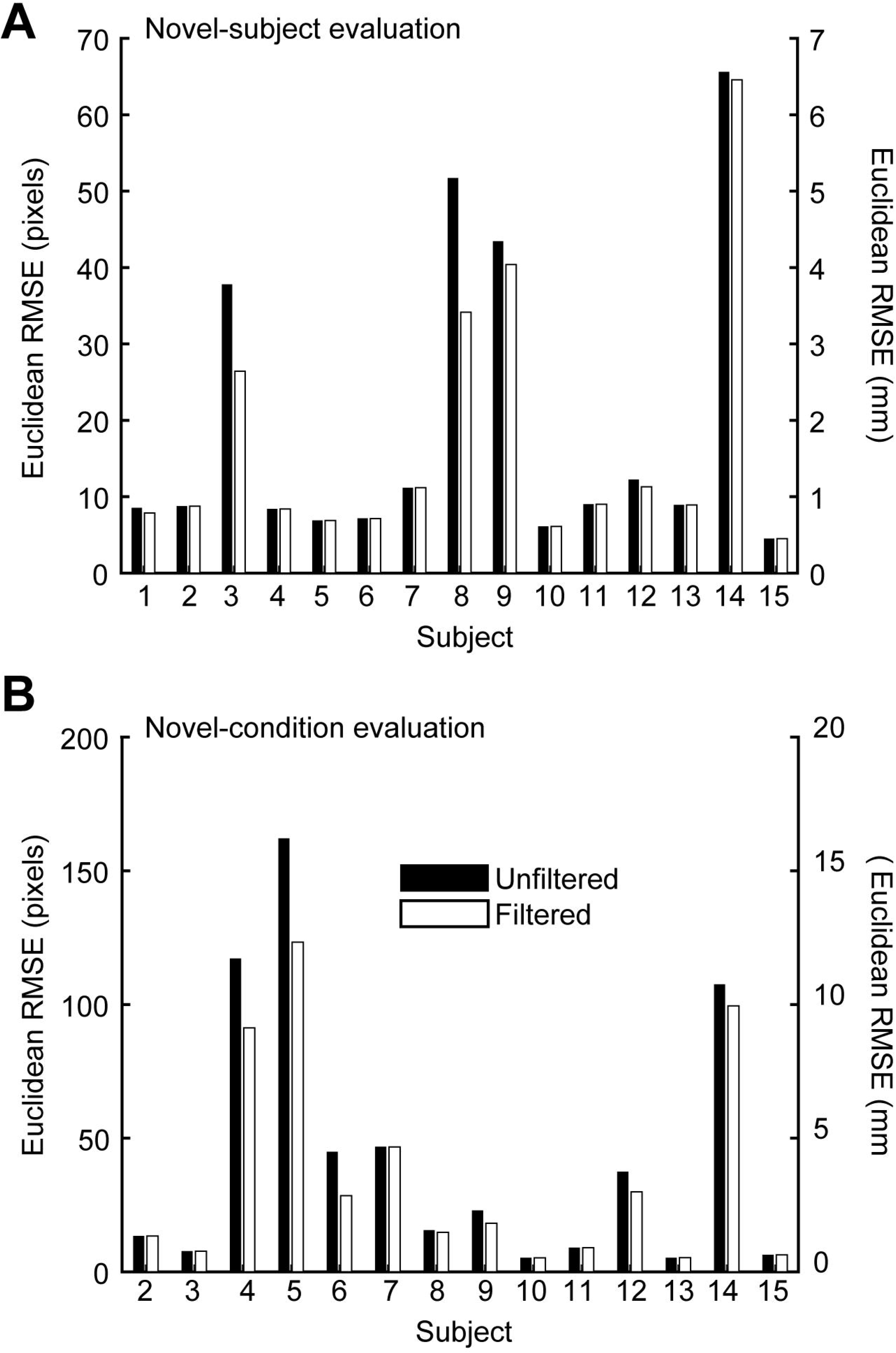
Unfiltered (black) and filtered (white) root mean square error (RMSE) for A) each individual subject in the novel subject evaluation (walking), and B) each individual subject in the novel condition evaluation (MVIC). RMSE is reported in pixels (left axis) and mm (right axis).

**Table 1.** Model performance metrics (mm)

| | RMSE | MAE (mean±SD) |
| --- | --- | --- |
| Novel-subject (walking) |  |  |
| Unfiltered | 2.72 | 1.26±1.30 |
| Filtered | 2.37 | 1.17±1.23 |
| Novel-condition (MVIC) |  |  |
| Unfiltered | 6.24 | 2.61±3.31 |
| Filtered | 5.05 | 2.18±2.64 |

RMSE: root mean square error, MAE: mean absolute error

### Novel-condition Evaluation: medial gastrocnemius MTJ displacements during MVICs

Overall unfiltered RMSE in the novel-condition evaluation was 6.23 mm, with an average unfiltered MAE of 2.61±3.31 mm (Table 1). The modified median filter reduced overall novel-condition RMSE by 1.18 mm (5.05 mm), and average MAE by 0.43 mm (2.18±2.64 mm) (Table 1). 88 and 90% of unfiltered and filtered predicted MTJ positions were classified as valid (Table 2). MAE for three subjects was greater than two standard deviations from the mean (Fig. 3B). Removing these subjects resulted in smaller overall RMSE (Unfiltered: 2.48 mm; Filtered: 2.10 mm) and smaller average MAE (Unfiltered: 1.05±0.83 mm; Filtered: 0.92±0.60 mm).

**Table 2.** Subject-specific and average prediction validity (%)

|  | Novel-subject (walking) |  | Novel-condition (MVC) |  |
| --- | --- | --- | --- | --- |
|  | Unfiltered | Filtered | Unfiltered | Filtered |
| Sub001 | 100 | 100 | - | - |
| Sub002 | 100 | 100 | 100 | 100 |
| Sub003 | 90 | 96 | 100 | 100 |
| Sub004 | 100 | 100 | 68 | 80 |
| Sub005 | 100 | 100 | 48 | 55 |
| Sub006 | 100 | 100 | 93 | 93 |
| Sub007 | 100 | 100 | 98 | 98 |
| Sub008 | 88 | 91 | 100 | 100 |
| Sub009 | 71 | 73 | 95 | 98 |
| Sub010 | 100 | 100 | 100 | 100 |
| Sub011 | 100 | 100 | 100 | 100 |
| Sub012 | 100 | 100 | 83 | 93 |
| Sub013 | 100 | 100 | 100 | 100 |
| Sub014 | 61 | 63 | 48 | 50 |
| Sub015 | 100 | 100 | 100 | 100 |
| Mean | 94 | 95 | 88 | 90 |

Values represent the proportion of predictions that were within 5 mm of the ground truth.

### Inference Speed

Inference speed was fastest on Nvidia GeForce GTX1080 (25.55±1.39 frames/s), followed by Nvidia Tesla V100-SXM2 (9.10±1.10 frames/s). CPU inference speed was considerably slower than GPU inference speed and was slowest with a smaller batch size (64 batch: 0.90±0.04 frames/s, 8 batch: 0.59±0.01 frames/s).

## Discussion

The purpose of this paper was to evaluate the efficacy of deep neural networks constructed with open-source software (15, 16) to accurately track medial gastrocnemius MTJ positions in cine B-mode ultrasound images collected during muscle actions spanning isolated contractions to walking. Traditional analysis of muscle-tendon kinematics using in vivo ultrasound imaging requires a time-consuming and labor-intensive process of manual labeling from one frame to the next. Here, we show that deep neural networks, trained using a small subject pool typical of biomechanics research studies, are effective and efficient for automated tracking of MTJ positions across a diverse range of contraction types. Accordingly, based on our cumulative findings, we feel confident advocating for the use of these deep neural networks as a suitable alternative to manual tracking of MTJ position from *in vivo* ultrasound images.

Novel-subject evaluation performance during walking was stronger than the performance of some semi-automated tracking techniques that use a combination of automatic and manual tracking methods (19, 20). Additionally, our networks outperformed those introduced recently by Leitner et al. (2020), who used deep learning methods to train and track MTJ positions during isolated contractions (14). Leitner et al. (2020) report a novel-subject MAE of 2.55 mm, with 88% of frames classified as valid using a 5 mm tolerance radius, compared to our novel-subject MAE of 1.26 mm with 94% of frames classified as valid. We note three subjects with outlier MAE values in the novel-subject evaluation (Fig. 3A). These outliers may be the result of our intentionally small subject pool to represent sample sizes common in biomechanics studies; our networks were not trained on an expansive catalog of MTJ characteristics, resulting in larger errors when presented with an uncharacteristic MTJ feature, such as narrow space between the deep and superficial aponeuroses (i.e. small medial gastrocnemius muscle width), or aponeuroses that don’t fully converge (see Supplemental Figure 1).

Overall RMSE was higher (i.e., worse performance) in the novel-condition evaluation than in the novel-subject evaluation. The relatively worse performance in the novel-condition evaluation may be due to differences in tissue deformation (i.e. muscle shape change) in maximal (MVICs) vs submaximal (walking) contractions. Thus, since our networks were only trained using images collected during walking, they may have been less capable of identifying MTJ image characteristics during maximal contractions. We purposefully withheld MVC images from the training set in order to assess the generalizability of the training set in a dynamic condition other than walking. We found that even when presented with novel-condition images, our networks performed with similar RMSE compared to semiautomated tracking techniques (20), and similar RMSE and MAE compared to fully automated deeplearning techniques (14)

Filtering the data with a modified median filter resulted in moderate reductions in overall RMSE and MAE for both the novel-subject and novel-condition evaluations. The filter was only applied to individual frames in which the confidence score of the predicted MTJ position was less than was <0.98. Subjects with MAE values greater than two standard deviations from the mean had a higher proportion of predictions with confidence scores <0.98 (Novel-subject evaluation: 28%, Novel-condition evaluation: 54%) compared to the remaining subject set(Novel-subject evaluation: 3%, Novel-condition evaluation: 13%). As such, when these subjects were removed, the difference between unfiltered and filtered values was relatively small in both the novel-subject and novel-condition evaluations. A modified median filter may be an ideal method for maintaining data integrity, while also smoothing erroneous data points.

We note a few limitations to this technique. First, the networks we describe here are only appropriate for tracking videos in which the MTJ is visible in all frames. Future studies should explore options to manage frames in which the MTJ is not visible, such as specifically training networks to infer position when the target is out of view (21). Additionally, our networks were based on MobileNetV2-1.0 (18), which is a shallower network (compared to ResNet network architectures) and may not be suitable for large images without downscaling. However, this network allows for faster training and analysis and can be run without a high-end GPU, which makes it ideal for researchers with limited computing resources. The fast processing speed of MobileNet also makes it ideal for future studies that aim to track in real-time. Finally, except for increasing the batch size, we used default DeepLabCut settings in order for these methods to be easily replicated. However, fine-tuning the settings may result in faster or more accurate performance.

There are many potential future directions for automated ultrasound image processing in the biomechanics and rehabilitation fields. For example, automated tracking with a trained network does not require the level of skill and experience of manual MTJ tracking. Thus, publicly available trained networks may encourage the adoption of tracking techniques in other fields, such as clinical diagnostics. Additionally, further development of these networks may facilitate real-time MTJ tracking, which could be a useful tool in rehabilitation and sports performance settings. Direct measurement of MTJ position from cine B-mode ultrasound images is important for reliably estimating tendon behavior and muscletendon interaction during functional activities. Although it is possible to indirectly measure tendon behavior by subtracting muscle fascicle length from muscle-tendon unit length, this indirect measurement frequently yields implausible outcomes (22). Thus, tools that can quickly and accurately estimate MTJ position have the potential to not only improve data processing speed, but also accelerate scientific discovery. Our results provide support for the use of open-source software for creating deep neural networks to reliably track MTJ positions in B-mode ultrasound images.

## Acknowledgements

This work was supported by a grant from the National Institutes of Health (R01AG058615). We also thank the University of North Carolina at Chapel Hill and the Research Computing group for providing computational resources and support.

