## Supplemental Figure 1 for "Automated Analysis of Medial Gastrocnemius Muscle-Tendon Junction Displacements During Isolated Contractions and Walking Using Deep Neural Networks"

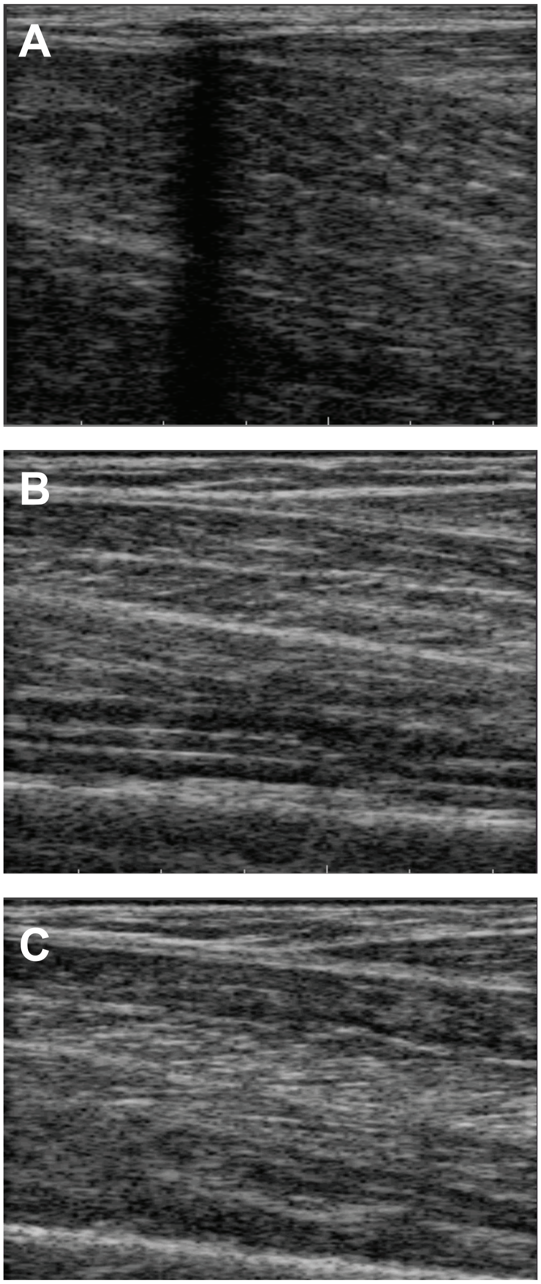


Supplemental Figure 1. Examples of image characteristics that produced poor tracking results. A) acoustic shadow, B) aponeuroses do not converge, C) more than one ‘Y’ shape on superficial aponeurosis
